# Cryo-EM structure of yeast 90S preribosome at 3.4 Å resolution

**DOI:** 10.1101/2020.09.16.299644

**Authors:** Yifei Du, Weidong An, Keqiong Ye

## Abstract

Previously, we determined a cryo-EM structure of Saccharomyces cerevisiae 90S preribosome obtained after depletion of the RNA helicase Mtr4 at 4.5 Å resolution (Sun et al., 2017). The 90S preribosome is an early assembly intermediate of small ribosomal subunit. Here, the structure was improved to 3.4 Å resolution and reveals many previously unresolved structures and interactions, in particular around the central domain of 18S rRNA. The central domain adopts a closed conformation in our structure, in contrast to an open conformation in another high-resolution structure of S. cerevisiae 90S. The new model of 90S would serve as a better reference for investigation of the assembly mechanism of small ribosomal subunit.

## Introduction

The ribosome is a large protein-synthesizing machine composed of a small 40S subunit (SSU) and a large 60S subunit (LSU). Assembly of the yeast S. cerevisiae ribosome from four ribosomal RNAs (18S, 25S, 5.8S and 5S rRNAs) and 79 ribosomal proteins (RPs) proceeds via a series of preribosomal particles and depends on the coordinated action of more than 200 trans-acting protein assembly factors (AFs) and many small nucleolar RNAs (snoRNAs) (Woolford and Baserga 2013; Bassler and Hurt 2019; Klinge and Woolford 2019). These factors mediate the modification, processing and folding of rRNAs, association of RPs and structural remodeling of preribosomes.

Ribosome assembly starts in the nucleolus with the transcription of a long 35S pre-rRNA that contains sequences for 18S, 5.8S and 25S rRNAs and four external and internal transcribed spacers (ETS and ITS). The 5’ region of 35S pre-rRNA, spanning the 5’ ETS, 18S and ITS1, is co-transcriptionally stepwise assembled into an early precursor of the SSU, known as the 90S preribosome or small subunit processome (Osheim et al. 2004; Chaker-Margot et al. 2015; Zhang et al. 2016). The 5’ ETS plays a critical role in 90S assembly by recruiting numerous AFs, including three independently formed subcomplexes: UTPA, UTPB and U3 snoRNP, to form a 5’ ETS particle (Chaker-Margot et al. 2015; Zhang et al. 2016; Hunziker et al. 2019). The 5’ and central domains of 18S each recruit several AFs independently of the 5’ ETS particle (Hunziker et al. 2019; Chen et al. 2020), whereas the 3’ major domain initially recruits few AFs. When the 18S rRNA is nearly completely transcribed, a dozen of late factors associate and the labile factors recruited by the 18S region, including U14 and snR30 snoRNA, dissociate. Consequently, the 90S compacts into a dense structure capable of processing the pre-rRNA. After the 5’ ETS is cleaved at the A0 and A1 sites and the ITS1 is cleaved at the A2 site, the 90S is transformed into a pre-40S that further maturates in the cytoplasm.

Several structures of 90S preribosome from S. cerevisiae and the thermophilic yeast Chaetomium thermophilum have been solved by single particle cryo-electron microscopy (cryo-EM) (Kornprobst et al. 2016; Barandun et al. 2017; Chaker-Margot et al. 2017; Cheng et al. 2017; Sun et al. 2017; Cheng et al. 2019). The highest resolution structure of S. cerevisiae 90S was determined at 3.8 Å (Barandun et al. 2017; Chaker-Margot et al. 2017). However, in this structure, the central domain was resolved at ∼7 Å and adopts an open conformation that greatly differs from a closed conformation in other 90S structures. The open conformation may represent a suppressed state since the 90S particle (called S-90S hereafter) was purified under starvation condition where ribosome assembly was inhibited. The structure of C. thermophilum 90S (Ct-90S) has been determined to 3.2-3.5 Å (Kornprobst et al. 2016; Cheng et al. 2017; Cheng et al. 2019). Multiple states of Ct-90S were observed, including major states with the central domain in the closed conformation and minor states with the central domain in the open conformation. The C. thermophilum and S. cerevisiae 90S are generally similar in structure and also display some differences. For example, their 5’ ETS are highly divergent in sequence. In Ct-90S, the assembly factor Rrt14 is missing and Utp9 in the UTPA complex is replaced by a second copy of Utp5 that lacks WD domain.

We previously determined cryo-EM structures for a wild type S. cerevisiae 90S and two mutant 90S with the RNA helicase Mtr4 or Dhr1 depleted. The two helicases were depleted in order to block the progression of 90S and improve the sample homogeneity. The structure of Mtr4-depleted 90S was determined at 4.5 Å, where the other two structures determined at ∼8 Å. These structures all have a closed conformation in the central domain. These structures were built mainly on homology models and contained errors as compared to the high-resolution structure of S-90S. Therefore, an accurate structure is needed for S. cerevisiae 90S in the closed conformation, which is more physiologically relevant than the open conformation. Moreover, because of the varied resolution over the large volume of 90S density maps, there are still many parts that are invisible or cannot be accurately modeled due to missing or poor density. Analysis of different 90S particles would be helpful to resolve the missing structures and understand progression of 90S.

Here, we have reanalyzed the Mtr4-depleted 90S by cryo-EM and built a model at 3.4 Å. The revised model reveals many new structures and interactions and enhances our understanding about the organization of the complicated RNA-protein complex.

## Results and Discussion

### Structure determination

To improve the model of Mtr4-depleted 90S, we recollected micrographs in a Titan Krios microscopy (Figure 1-figure supplement 1). Compared with our previous analysis, the new dataset contained more images (4714 vs 1102) and were collected on a different camera (K2 Summit vs Falcon III) and with a smaller pixel size (1.06 Å vs 1.76 Å). In addition, an iterative particle-picking strategy was developed to retrieve more particles from micrographs (See Materials and Methods). A single high-resolution state was sorted out in 3-dimensional (3D) classification, indicating that the sample was highly homogeneous. A density map was reconstructed from 128,637 particles at an overall resolution of 3.4 Å (Figure 1A, Figure 1-figure supplement 1). Focused classification and refinement were conducted to improve local density for the core region, the 5’ domain, the central domain, the 3’ major domain and the UTPA complex (Figure 1-figure supplement 1C and 2). Side chains of protein and bases of RNA were discernible at the well resolved regions (Figure 1-figure supplement 3). The new model was built on the overall and focused maps using an initial model that combines the previously determined structures of S. cerevisiae 90S (Barandun et al. 2017; Sun et al. 2017) (Supplementary Table 1).

**Figure 1.**
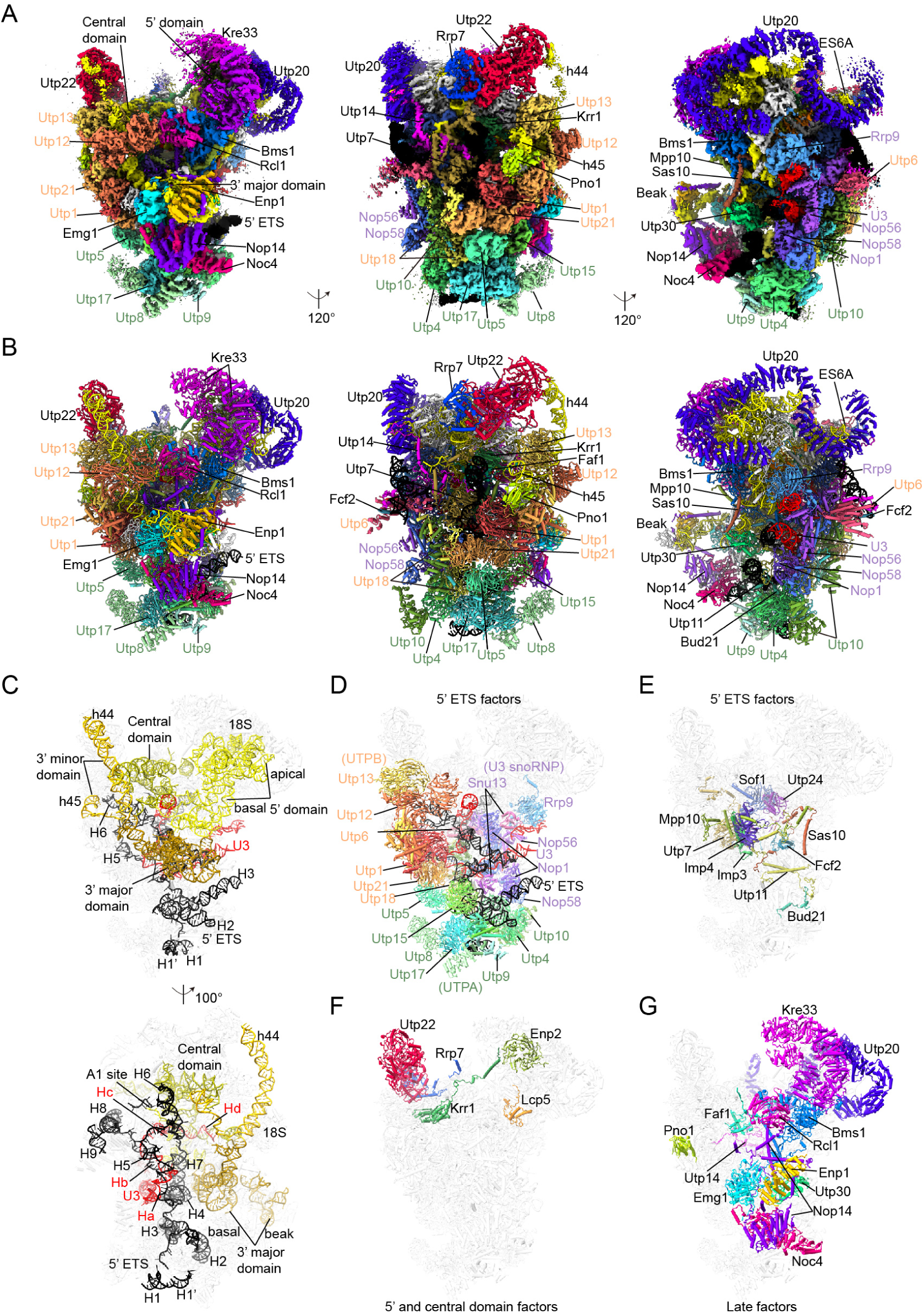
Cryo-EM structure of Mtr4-depleted 90S preribosome. (A) Composite cryo-EM maps of Mtr4-depleted 90S shown in three views. The overall map and five focused maps are overlaid. 5’ ETS, 18S and U3 RNAs are colored in black, yellow and red. Individual AFs are colored differently, all RPs are white and the unmodeled densities are grey. The UTPA, UTPB and U3 snoRNP components are labeled in green, orange and purple, respectively. (B) Cartoon representation of Mtr4-depleted 90S structure. (C) Structure of 5’ ETS, 18S and U3 RNAs shown in two views. The other components of 90S are displayed as transparent background. (D-G) Illustration of AFs by their assembly order. The UTPA, UTPB and U3 snoRNP complexes (D) and other AFs (E) recruited by the 5’ ETS, the factors recruited by 5’ and central domains of 18S rRNA (F), and the lastly assembled late factors (D) are shown on transparent background of 90S structure. The structures have the same orientation in the left panels of A and B, the upper panel of C and all panels of D-G.

The new model consists of 5’ ETS, 18S and U3 RNAs, 18 RPs and 49 AFs (Figure 1A-B and Supplementary Table 2). The model of Krr1, Faf1, Rrp7 and Utp7 were significantly extended. An atomic model has been built for the tandem WD domains of Utp13 that previously was a poly-Ala model. Rrt14 and Rrp5 were not included due to their extremely weak density. Rrt14 is a non-essential AF present in some yeasts, but not in C. thermophilum (Hontz et al. 2009; Chaker-Margot et al. 2015; Zhang et al. 2016). In the S-90S structure, a middle part of Rrt14 contacts Utp11 and the 5’ ETS and a C-terminal part of Rrt14 binds Utp30 (Barandun et al. 2017) (Figure 1-figure supplement 5A-B). The densities of both parts were absent in our map (Figure 1-figure supplement 5C), suggesting that Rrt14 is not bound to the two sites in Mtr4-depleted 90S. The TPR domain of Rrp5 that originally binds Utp22 was not modeled either due to extremely weak density (Figure 1-figure supplement 5D).

### Overall structure

Mtr4-depleted 90S adopts the same globular shape as its previously determined structure and other 90S structures (Figure 1A-B). The 5’ ETS, 18S and U3 RNA are largely buried inside and coated by abundant proteins. The 5’ ETS RNA folds into a highly branched structure and assembles with abundant AFs into the 5’ ETS particle that constitutes the bulk of the structure body (Figure 1C-E). The 5’ domain and the central domain of 18S rRNA with associated RPs and AFs are placed on the top of the structure and the 3’ major domain is located at the center (Figure 1C). Helices 44 and 45 in the 3’ minor domain have not yet packed into the 5’ and central domain, respectively. Overall, the four domains of 18S rRNA are partially assembled and segregated without forming the global architecture of the mature SSU (Figure 1C, Figure1-figure supplement 4). The 5’ ETS and 18S rRNA are linked by the 5’ domain of U3 snoRNA that form helices Ha and Hb with the 5’ ETS and helices Hc and Hd with the 18S rRNA.

The A1 site junction between the 5’ ETS and 18S rRNA is intact (Figure 1C, Figure 1-figure supplement 3N), indicating that the structure represents a state before cleavage of A1 site, similar with the S-90S and Ct-90S structures.

The three prominent complexes UTPA, UTPB and U3 snoRNP, that bind the 5’ ETS at the earliest stage of 90S assembly, constitute the base and the back face of the structure (Figure 1D) (relative to the view on the left panel of Figure 1A). The other 5’ ETS factors are mostly located inside the structure (Figure 1E). The factors initially recruited by the 5’ and central domain bind to these domains on the top of the structure (Figure 1F). The late factors that join 90S at the last stage of assembly are mostly situated at the front face and connect the ribosomal domains and the 5’ ETS particle into a compact structure (Figure 1G).

A large portion of the structure of Mtr4-depleted 90S is consistent with the previously determined high-resolution structure of S-90S (Barandun et al. 2017). Major differences lie in the central domain that adopts an open conformation and poorly resolved (∼ 7 Å) in S-90S and a closed conformation in Mtr4-depeleted 90S (Figure 2A, C). The central domain was resolved at 3.6 Å in our maps, which allowed for de novo modeling. The central domain also adopts the closed state in major states of Ct-90S (Figure 2B), but is less resolved compared to our structure.

**Figure 2.**
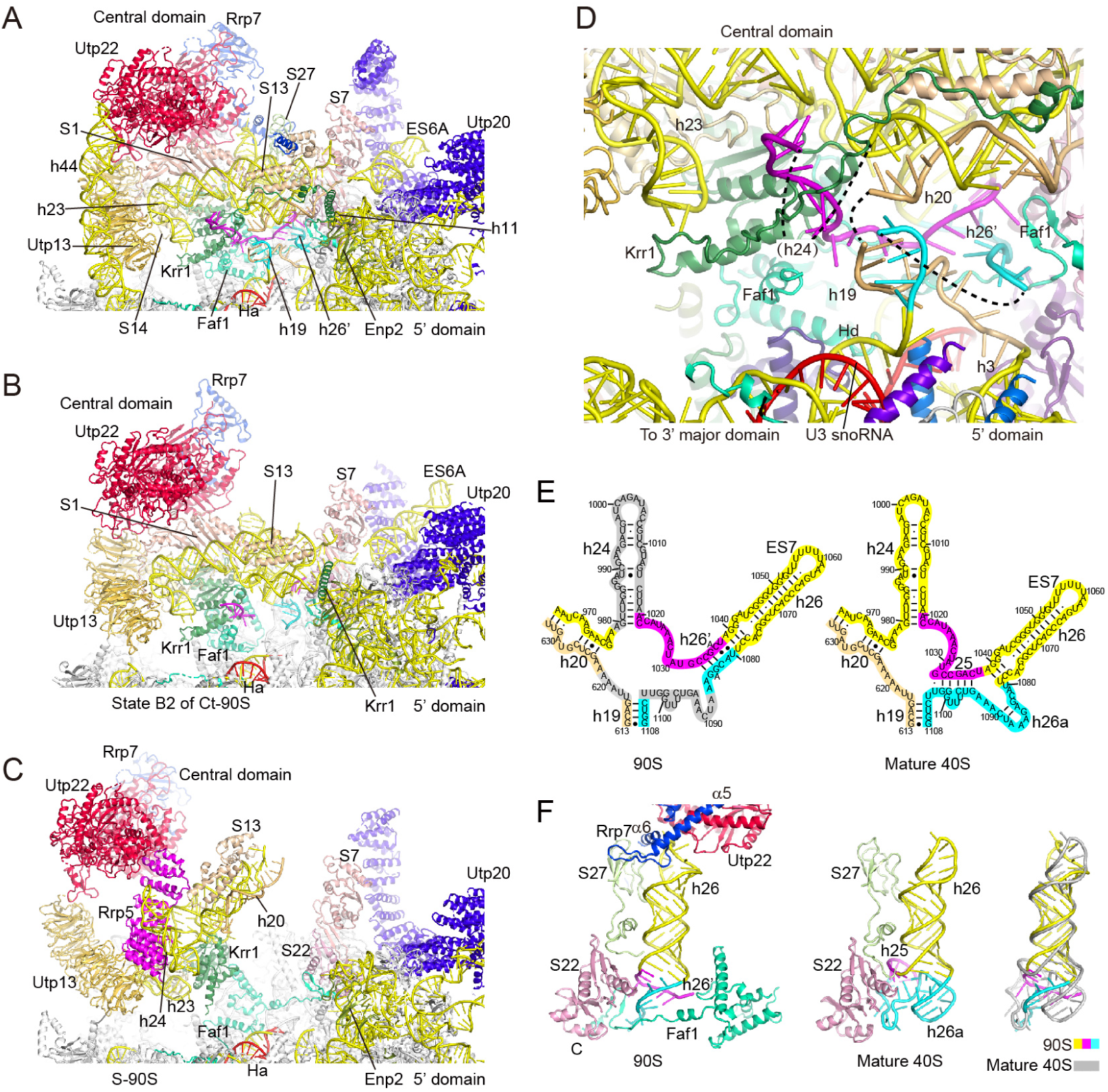
Structure of the central domain. (A-C) Structure of the central domain in Mtr4-depleted 90S (A), state B2 of Ct-90S (PDB code: 6RXV), S-90S (B) and S-90S (5WLC) (C). (D) Interface of the central domain to the 5’ domain and 3’ major domain. Segments of 18S rRNA are colored as in E. The dashed lines represent disordered RNA regions. (E) Secondary structure diagrams of 18S rRNA around helix 26 in 90S and mature 40S ribosomes. RNA segments are shaded in different colors. The unmodelled regions in the 90S are shaded in grey. (F) Structure of helix 26 in 90S (left) and mature 40S (middle) ribosomes and their alignment (right).

### The central domain

The platform in the mature SSU is made up by the bulk of the central domain of 18S rRNA (helices 19-27), helix 45 from the 3’ minor domain, and seven RPs (Rps1, Rps7, Rps13, Rps14, Rps22, Rps26 and Rps27). In the 90S, the platform is partially assembled from helices 20, 22, 23 and 26 and all seven RPs but Rps26, and bound by assembly factors Krr1, Faf1, Utp13, Utp22, Rrp7 and Utp20 (Figure 2A). Importantly, Krr1 occupies the position of helix 45 and Rps26, excluding their incorporation.

Extension segment 6 (ES6) is a large eukaryote specific insertion in the central domain involved in binding the 5’ domain. Helix A of ES6 (ES6A) is partially resolved in the new model. ES6A sticks out of the central domain and is encapsulated by the C-terminal half of the long super-helical structure of Utp20 (Figure 2A).

The interface of the central domain to other ribosomal domains was poorly resolved in all previously determined 90S structures (Figure 2B-C). Located at the center of the interface is helix 19 that covalently connects the immature platform to the 5’ domain and the Hd helix that leads to the 3’ major domain (Figure 1-figure supplement 4). Helix 19 and its linker to helices Hd and h3 are structured, but it linkers to helices 20 and 26 are disordered in the new 90S model (Figure 2D). Non-native RNA structures are formed in the domain interface. The short helices 25 and 26a in the mature SSU are not formed, but part of their sequences combine into a new 5-bp stem, named helix 26’, extending the base of helix 26 (Figure 2D-F). The formation of helix 26’ is stabilized by Faf1, ribosomal protein Rps22 and the linker RNA between helices 19 and 3.

### Utp22 and Rrp7

Utp22 and Rrp7 form a tight UTPC complex that binds at the top of the central domain. Their previous models in 90S are based on a crystal structure of their complex and lack the ∼100-residue, conserved and functionally important C-terminal domain (CTD) of Rrp7 (Lin et al. 2013). The CTD of Rrp7 and new interactions of Utp22 were visualized in the new model.

The resolved CTD of Rrp7 contains three helices α5-α7 and a few loops and binds at the top of the central domain (Figures 3A-D). Helix α6 and its N-terminal loop bind at the tip of helix 26 from the major groove side, whereas Utp22 binds the RNA from the minor groove side (Figure 3C). The two proteins mainly contact the phosphate and sugar groups of the RNA and also contact the base edges of U1058 and U1056 by potential sequence-specific interactions. The direct interaction of Rrp7 with 18S rRNA accounts for their crosslinking (Lin et al. 2013). The conformation of the RNA-binding loop of Rrp7 is further stabilized by interaction with Rps27, including an anti-parallel β-strand pairing (Figure 3C). The binding of Rrp7 shifts the tip of helix 26 from its mature position by ∼19 Å, but does not affect the position of Rps27 that mainly contacts the stem of helix 26 (Figure 2F).

**Figure 3.**
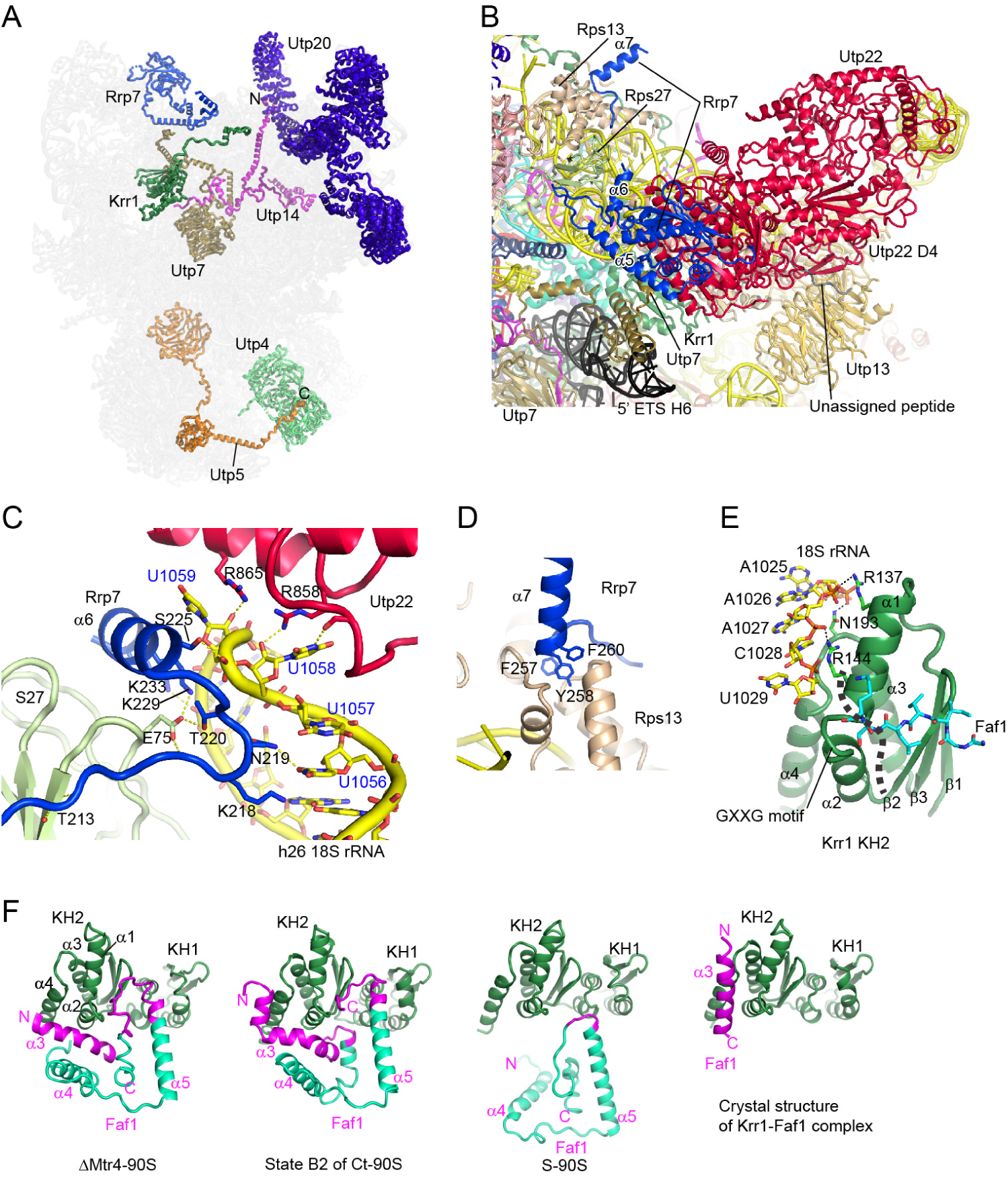
Expended interactions in the new model of 90S. (A) Rrp7, Utp7, Krr1, Utp14 and Utp5 contain extended tails. The other parts of 90S are shown as transparent background. (B) Newly resolved interactions of Utp22 and Rrp7. (C) The tip of 18S helix 26 is sandwiched between Utp22 and Rrp7. (D) Interaction of Rrp7 with Rps13. (E) The KH2 domain of Krr1 binds a single-stranded region of 18S rRNA with an atypical interface. The classic RNA-binding groove is marked by a dashed line. (F) Interaction between Faf1 and Krr1 in Mtr4-depleted 90S, state B2 of Ct-90S (PDB code: 6RXV), S-90S (5WLC) and a crystal structure of their complex (4QMF). The Krr1-binding regions of Faf1 are colored in magenta.

The N-terminus of the α7 helix of Rrp7 docks on Rps13 by inserting three phenyl rings into a hydrophobic pocket of Rps13 (Figure 3D). These aromatic residues are highly conserved (Lin et al. 2013), underscoring the importance of the interaction. Helix α7 is followed by highly conserved 25 amino acid residues that fold into an α-helix as shown by NMR analysis (Lin et al. 2013), but are missing in our structure. These residues likely mediate important interactions in other states of 90S.

Our model shows that Utp22 binds two additional proteins (Figure 3B). The C-terminal residues of Utp7 dock on a composite pocket formed by Utp22 and Rrp7. In addition, the D4 domain of Utp22 accepts a short peptide from an unassigned protein (Figure 3B).

### Krr1 and Faf1

Krr1 is composed of two RNA-binding K homology (KH) domains (KH1 and KH2) and a long C-terminal tail (Zheng et al. 2014). We found that a single-stranded RNA located 5’ of helix 26’ is sandwiched between the KH2 domain and the C-terminal tail of Krr1 (Figure 2D). The KH2 domain possesses a classic RNA-binding surface of KH domain, as indicated by the conserved GXXG motif between helices α1 and α2. However, the RNA binds at helix α1 and the loop connecting the α3 and α4 helix of the KH2 domain, rather at the classic RNA-binding groove, which is occluded by Faf1 (Figure 3E). The RNA binding is mainly directed to its phosphate groups.

Krr1 contains a ∼100-residue C-terminal tail that travels over a long distance from the central domain to the 5’ domain (Figure 2A). The tail interacts with Rps13 and helix 20 in the central domain and Enp2 and helix 11 in the 5’ domain. A similar interaction of the tail with the 5’ domain was also present in the Ct-90S structure (Figure 2B).

Faf1 is a binding partner of Krr1 (Karkusiewicz et al. 2004; Zheng et al. 2014). The helix α3 and the loop downstream of helix α5 of Faf1 bind Krr1 in a similar manner as in state B2 of Ct-90S (Cheng et al., 2019) (Figure 3F). Of note, helix α3 of Faf1 binds differently with Krr1 in a crystal structure of their complex (Zheng et al. 2014). Helix α3 of Faf1 is perpendicular to helix α4 of Krr1 in the 90S structure, but nearly parallel to helix α4 in the crystal structure. The crystal structure likely presents the strongest interaction mode between helix α3 of Faf1and Krr1 and could occur when they make the first contact during 90S assembly. Faf1 is assembled into 90S later than Krr1 and has the chance to meet Krr1 when 90S has not yet packed (Chaker-Margot et al. 2015; Zhang et al. 2016). Faf1 is essentially dissociated from Krr1 in the open conformation of the central domain as shown in the S-90S structure (Figure 2C, 3F). These structures suggest that the interface between Krr1 and Faf1 is plastic and prone to change during structural remodeling of 90S.

### New interactions involving the extended tails of Utp7, Utp14 and Utp5

Utp7 is an AF recruited by the 5’ ETS and its WD domain is bound at the body of 90S (Figure 3A). The new model of 90S shows that Utp7 contains an extended C-terminal tail that approaches the central domain and contacts with the 5’ ETS RNA (see below), Krr1, Utp22 and Rrp7 (Figure 3B).

Utp14 adopts a highly stretched structure and primarily contacts with the 5’ ETS particle (Figure 3A). The new model reveals that the N-terminal region of Utp14 attaches to the convex surface of the super-helical protein Utp20 (Figure 1-supplement 3O).

Utp5 is a component of the UTPA complex and locates at the base of the 90S structure. Utp5 was shown to make a two-hybrid interaction with Utp4 (Freed and Baserga 2010), but this interaction was not observed in the previous 90S structures. Our map showed that a C-terminal peptide of Utp5 binds the WD domain of Utp4 (Figure 3A, Figure 1-figure supplement 3P), accounting for the biochemical data.

### The 3’ minor domain of 18S rRNA

The 3’ minor domain of 18S rRNA is composed of helices 44 and 45 that are part of the body and platform in the mature SSU, respectively. Both helices are not yet packed in the 90S. The long helix 44 forms an arc linking the major domain of 18S rRNA to Utp22. The structure of h44 was not correctly registered with RNA sequence in the previous Mtr4-depleted 90S structure and largely missing in other structures (Figure 2B-C). The linker between h44 and h45 were also missing in previous 90S models. The full length of RNA from h44 to h45 has been modeled (Figure 4A-B). The entire length of h44 fully accounted for the long curved density linking Utp12 and Utp22. The linker between h44 and h45 adopts an extended conformation and sits on the C-terminal domain (CTD) tetramer of the UTPB complex.

**Figure 4.**
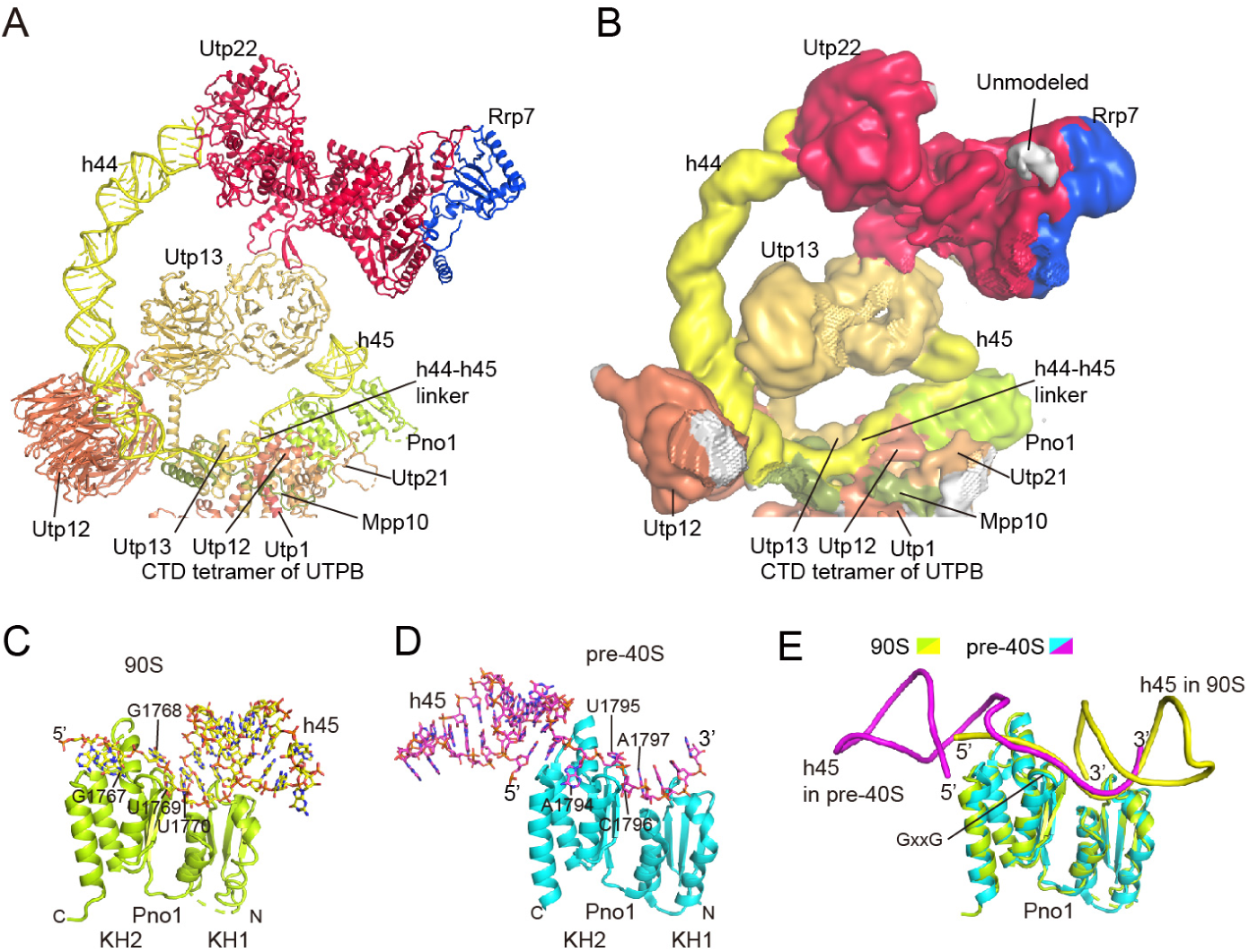
Structure of the 3’ minor domain of 18S rRNA. (A) Structure and interaction of the 3’ minor domain in Mtr4-depleted 90S. (B) Cryo-EM density map low-pass filtered to 10 Å for the same region of A. (C-D) RNA binding of Pno1 in the 90S (C) and pre-40S structure (PDB code: 6FAI) (D). (E) Aligned structures of the Pno1-RNA complex in 90S and pre-40S.

Pno1 contains two KH domains and is present in both 90S and pre-40S ribosomes (Schafer et al. 2003). In the 90S, Pno1 binds h45 with the degenerated KH1 domain and the single-stranded sequence 5’ of h45 with the KH2 domain that has a classic RNA-binding surface (Figure 4C). In the pre-40S ribosome (Heuer et al. 2017; Scaiola et al. 2018), Pno1 recognizes a different single-stranded sequence 3’ of h45. Two 4-nt RNAs are recognized by the Pno1 KH2 domain in both 90S and pre-40S, but their sequences are completely different, namely 1767-GGUU-1770 in 90S and 1794-AUCA-1797 in pre-40S. The KH1 domain, which lacks the GXXG motif, binds a duplex RNA in 90S and a single-stranded RNA in pre-40S. Although the target RNAs have different sequences and secondary structures, the RNA strands that directly contact Pno1 are superimposable (Figure 4E). Therefore, Pno1 employs the same surface to bind two RNA targets at different stages of ribosome maturation.

### A revised model of the 5’ ETS

The 5’ ETS is composed of ten helices (H1 to H10) and forms two intermolecular duplexes with U3 snoRNA (Ha and Hb) (Figure 5). The structure of H1 and the upper stem of H6 were revised or newly built. We will also discuss the structure of 5’ ETS in relation with its sequence conservation in the Saccharomycotina subphylum of yeast (Figure 5-figure supplement 1)(Chen et al. 2020).

**Figure 5.**
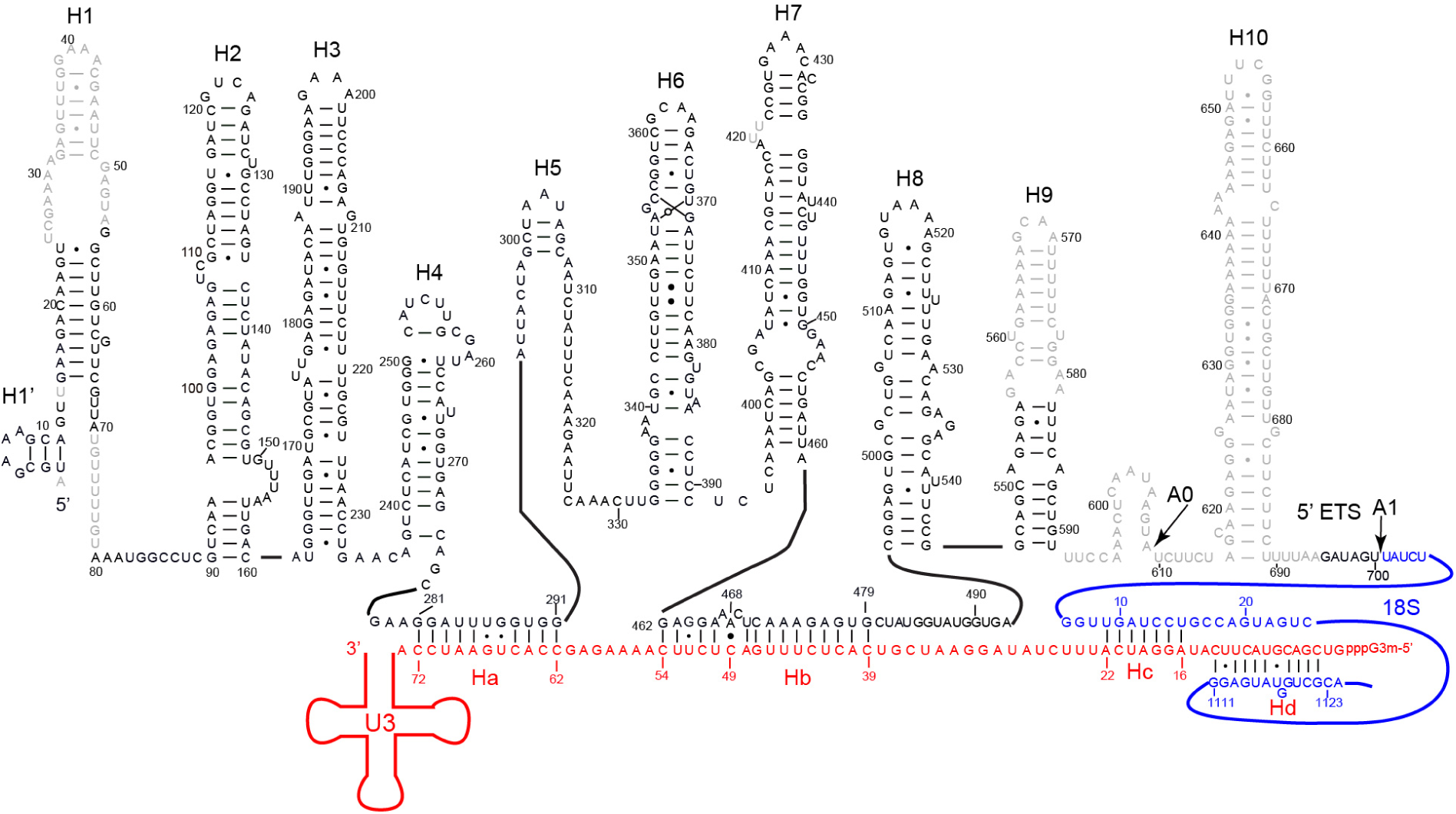
Secondary structural model of 5’ ETS. The model is based on the new structure of Mtr4-depleted 90S. The 5’ ETS, 18S and U3 RNAs are colored in black, blue and red, respectively. The unmodeled nucleotides are colored in grey and their secondary structures are predicted. Helixes of 5’ ETS are termed as H1 to H10 and the newly identified short helix at the 5’ end is named as H1’. The helices between U3 and 5’ ETS/18S RNA are named as Ha to Hd.

We found that the 5’ end sequence of the 5’ ETS folds into a 3-bp hairpin, termed helix H1’, rather than being part of helix H1 in the previous 90S models (Figure 5, 6A, Figure 1-figure supplement 3M). Helix H1’ is co-axially stacked on the shortened H1 helix. The sequence of H1’ is much more conserved than that of H1 (Figure 5-figure supplement 1), supporting that this region forms a separate structural unit. The high degree of conservation of H1’ in Saccharomycotina is consistent with its intimate interaction with the WD domain of Utp17 and the CTD of Utp9 (Figure 6A). Interestingly, the entire H1’-H1 region is not required for the processing of a pre-18S rRNA construct (Chen et al. 2020). The 5’ region of the C. thermophilum 5’ ETS is also made up of two co-axially stacked hairpins, which however differ from these of S. cerevisiae 5’ ETS in sequence, length and structural orientation in 90S (Figure 6B-C). The structure and interaction of the first two hairpins of 5’ ETS are rather variable in the two yeasts, which tends to agree with their minor role in 90S assembly.

**Figure 6.**
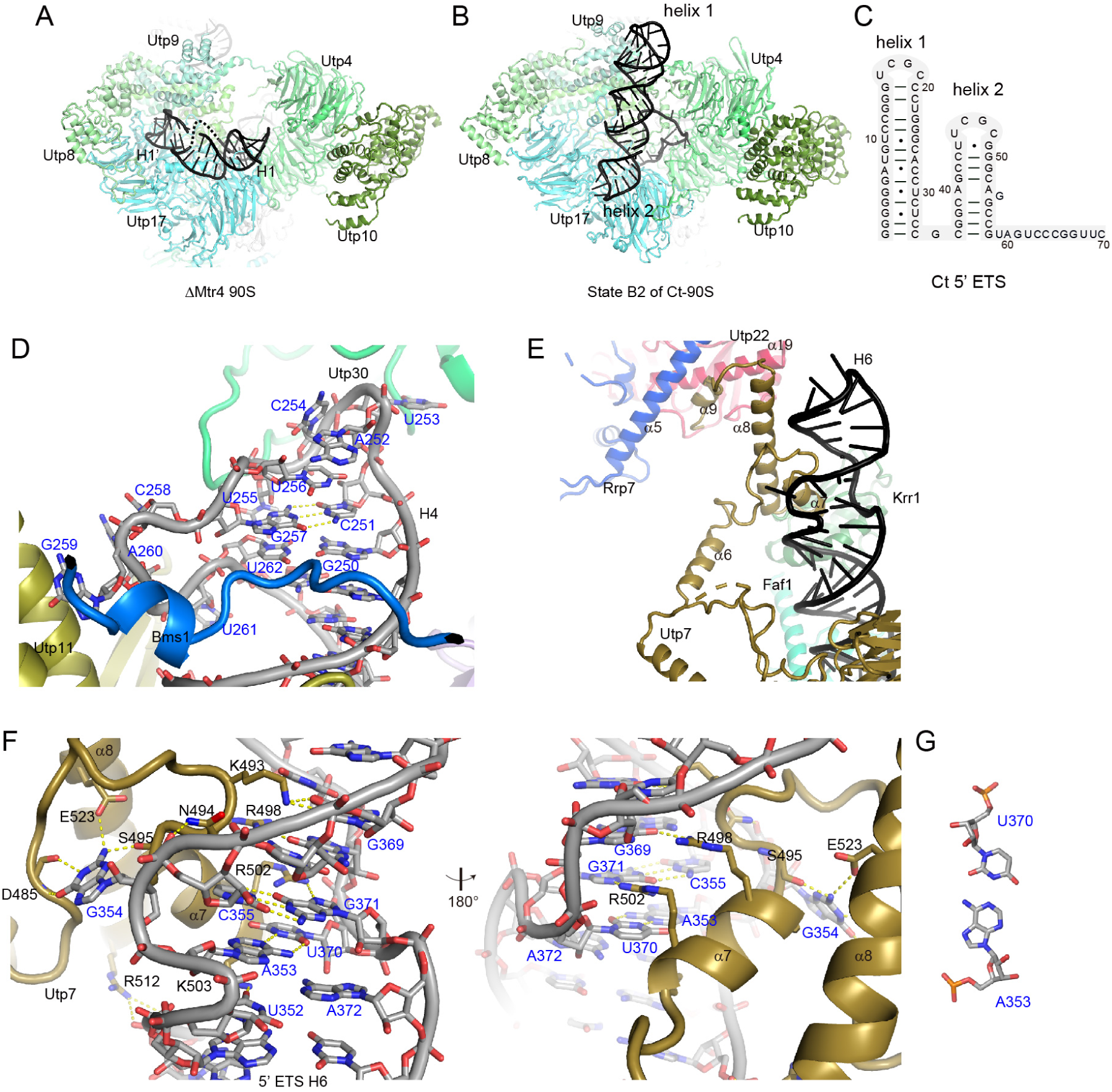
Structure and interaction of the 5’ ETS. (A) Helices H1’ and H1 of 5’ ETS in the Mtr4-depleted 90S structure. (B) Helices 1 and 2 of the C. thermophilum 5’ ETS adopt a different orientation in 90S structure (PDB code: 6RXV). (C) Secondary structure diagram of helix 1 and 2 of the Ct 5’ ETS. (D) Structure and interaction of the top loop of H4. (E) Structure and interaction of H6. (E) Detailed interaction between Utp7 and H6 shown in two opposite views. (G) A reverse Watson-Crick pair in H6.

The H4 helix of 5’ ETS is capped with a highly conserved 11-nt loop that interacts with Utp30, Utp11 and Bms1 in the 90S structure. Within the loop, C251 forms a base pair with G257 (Barandun et al. 2017). The two positions display strong co-variation in the 5’ ETS sequences from Saccharomycotina (Chen et al. 2020) (Figure 5-figure supplement 1), supporting their important role in forming the loop structure.

The new map shows that the upper stem of H6 forms a special structural motif that is bound by a two-helix domain in the C-terminal tail of Utp7 (Figure 6E-F). G354 on the 5’ strand of H6 bulges out and inserts into a pocket of Utp7. The base edge of G354 is specified by the carbonyl and amide groups of D485 and the side chains of E523 and S485. The phosphate backbone of the right strand of H6 makes an S-shaped twist at U370 and G371. As a result, the two stands of H6 pair in a parallel manner at the twist. A353 and U370 form a reverse Watson-Crick pair and C355 and G371 form a Watson-Crick pair. The α7 helix of Utp7 and its N-terminal loop insert into the major groove of H6 and contact both the base edges and backbone of the RNA. In addition, the α8 helix of Utp7 interacts with the phosphate backbone of H6 (Figure 6E). Utp7 appears to recognize the specific structure and sequence on the upper stem of H6. Consistently, the middle stem of H6 contacted by Utp7 is highly conserved in Saccharomycotina (Figure 5-figure supplement 1).

## Conclusion

We have built a more complete and accurate model for a fully assembly 90S preribosome. The model reveals new structures and interactions, in particular around the central domain, and expands the complicate protein-protein and protein-RNA interaction networks in the 90S. The central domain adopts a closed conformation in our structure, which is considered to be more physiologically relevant than the open conformation. The new 90S model would serve as a better reference for investigating the assembly mechanism of small ribosomal subunit.

## Materials and Methods

### Cryo-EM sample preparation and data collection

Mtr4-depleted 90S particles were purified from the yeast strain GAL::mtr4/Enp1-TAP (BY4741, Enp1-TAP::HisMX, natNT2-pGALL::3HA-mtr4) grown in YPD medium for 16 h, as previously described (Sun et al. 2017).

Quantifoil grids (1.2/1.3, 300 mesh) were covered with a thin layer of home-made continuous carbon film and glow-discharged in a Solarus 950 plasma cleaner (Gatan) for 40 sec. Cryo-EM grids were prepared in an FEI Vitrobot Mark IV. Aliquots of sample (3.5 μl, OD280 ∼2) were applied on grids at 4 °C and 100% of humidity for 30 sec. Grids were blotted for 3 sec with a force level of -3 and plunged into liquid ethane.

A total of 4,723 micrographs were collected on a 300 kV Titan Krios microscope (FEI) equipped with a K2 Summit camera (Gatan). Micrographs were recorded in the super-resolution mode with a pixel size of 0.53 Å, an exposure time of 9 sec and a total electron dose of 50 e^-^/Å^2^. Defocus values varied from 1.5 to 2.5 μm.

### Image processing

Micrographs composed of 32 frames were processed with MotionCor2 for frame alignment, gain correction and dose weighting (Zheng et al. 2017). The summed images were binned by a factor of 2, resulting in a pixel size of 1.06 Å. Magnification distortion was corrected by mag_distortion_correct_1.0.1 (Grant and Grigorieff 2015). Images without dose weighting were used for contrast transfer function (CTF) determination by CTFFIND 4.1.8 (Rohou and Grigorieff 2015). After CTF determination, images with poor thong rings were discarded. Dose weighted images were used for 3D reconstruction. Unless specifically mentioned, all following processing steps were performed using Relion (Scheres 2012; Kimanius et al. 2016).

Particles were auto-picked in Relion using 2D class averages filtered to 40 Å as template. The initial templates were generated from about 2000 manually picked particles. Particles were extracted using a box of 512 pixels (542 Å) in size. Background area was defined by a circle of 444 pixels (466 Å) in diameter. Particles were 3D classified into 5 classes using the EMD-6695 map low-pass filtered to 60 Å as an initial map.

An iterative particle-picking procedure was applied to increase the number of particles extracted from micrograph. Particles belonging to high-resolution classes from the last round of 3D classification were subjected to 2D classification. The resulting class averages were used as templates for the next round of auto-picking. After 3D classification of newly picked particles, particles in high-resolution classes were selected. The procedure was repeated for three times. All particles in high-resolution classes from three rounds of particle picking and 3D classification were merged and the overlapping particles were removed using an in-house written script. The non-redundant set of particles were subjected to another round of 3D classification and selection.

Datasets I (1978 images) and II (2736 images) were processed separately. Following three rounds of iterative particle-picking and 3D classification, 53,674 and 74,963 particles in high-resolution classes were obtained from datasets I and II, respectively. These particles all belonged to classes with similar structural features and were combined to reconstruct an overall density map at 3.4 Å, following 3D auto-refinement, CTF refinement and post-processing.

To improve density in region of interest, focused maps were calculated for the core region, the 5’ domain, the central domain, the 3’ major domain and the UTPA complex (Figure 1-figure supplement 2) (Bai et al. 2015; von Loeffelholz et al. 2017). The overall map was covered with a mask created in RELION. Particles were 3D classified by two rounds according to the masked volume. In the focused classification, particles were realigned for the core region map, but not for other focused maps. Particles from high-resolution classes were selected, subtracted of signals outside the mask and reconstructed. Resolution of map was estimated according to the gold-standard Fourier shell correlation=0.143 criteria (Scheres and Chen 2012). Local resolution was calculated by Resmap (Kucukelbir et al. 2014).

### Model building and refinement

An initial model was generated from previous determined structures of S. cerevisiae 90S (PDB code: 5WYK and 5WLC) (Barandun et al. 2017; Sun et al. 2017) and fitted into the overall map. The model was built in Coot (Emsley and Cowtan 2004), based on the overall and five focused maps. An initial homolog model for the tandem WD40 domains of Utp13 was built by SWISS-MODEL using the Gemin5 structure as template (PDB code: 5TEE) (Waterhouse et al. 2018).

The model was refined by phenix.real_space_refine with secondary structure and geometry restraints and individual B-factor determination (Adams et al. 2010). The model was refined against individual focused maps at the early stage of model building and against the overall map at the final stage. Statistics of data collection, structural refinement and model validation are listed in Supplementary Table S1. Composition of the model is listed in Supplementary Table S2. Structural figures were prepared with Chimera (Pettersen et al. 2004), ChimeraX (Goddard et al. 2018) and PyMOL (DeLano 2002).

### Database

The cryo-EM map and coordinates have been deposited in the EMDB and PDB with accession codes: EMD-9964 and 6KE6 (already released).

## Acknowledgments

Cryo-EM data were collected at the Center for Biological Imaging (CBI), Institute of Biophysics, Chinese Academy of Science (CAS). We thank Fei Sun, Xiaojun Huang, Zhenxi Guo, Gang Ji, Bolin Zhu and Deyin Fan at CBI for assistance in cryo-EM data collection and Xiuling Gao for technical assistance. The study was supported by National Natural Science Foundation of China [91940302, 91540201, 31430024], Strategic Priority Research Program of Chinese Academy of Sciences [XDB37010201] and National Key R&D Program of China [2017YFA0504600]. Author contributions: K.Y. initiated the project; Y.D. collected cryo-EM data and determined the structure; W.A. prepared samples, Y.D. and K.Y. analyzed the structure and wrote the paper.

---

**Figure 1-figure supplement 1.**
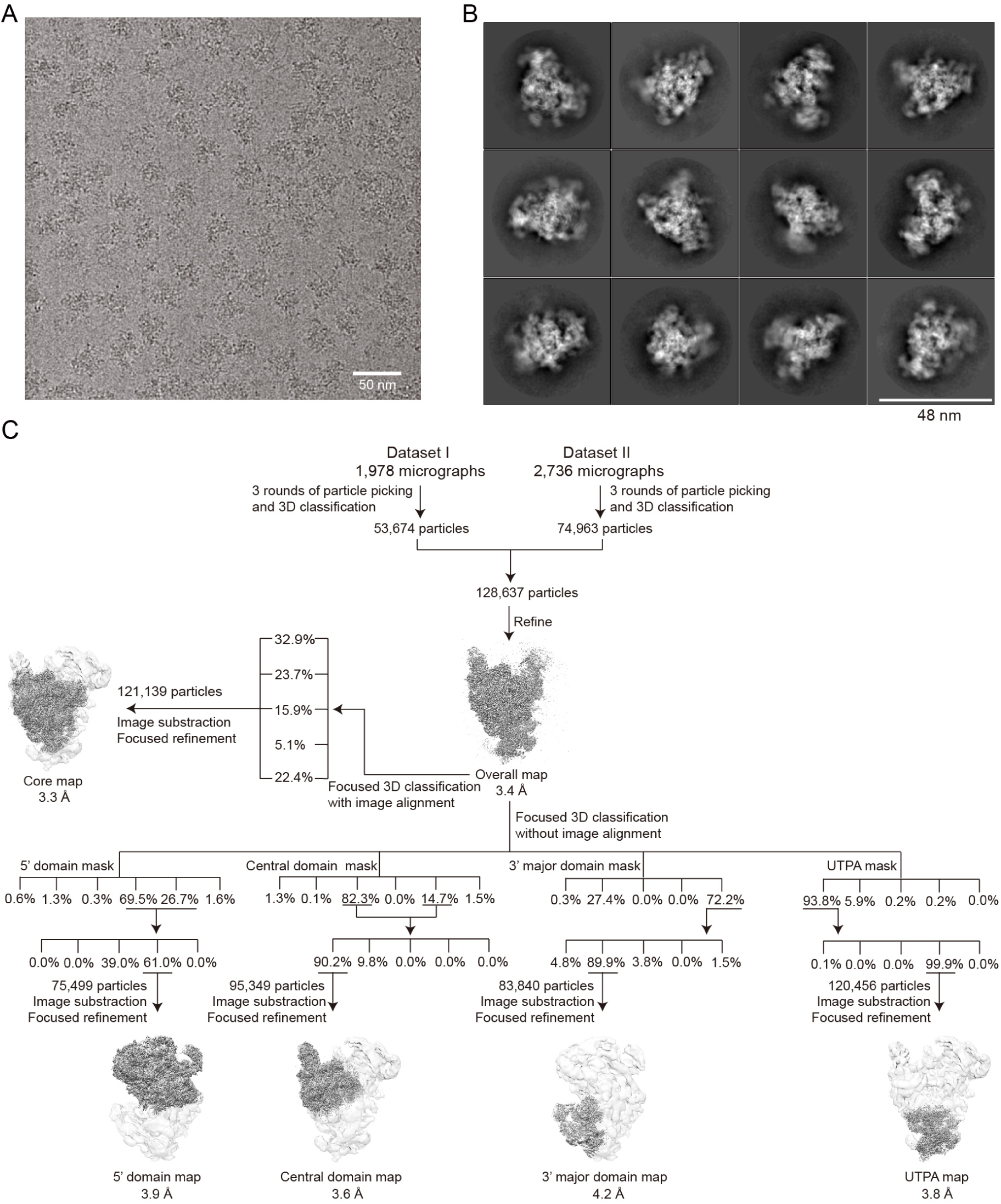
Cryo-EM analysis of Mtr4-depleted 90S. (A) A representative cryo-EM micrograph. Scale bar = 50 nm. (B) Selected 2D class averages. Scale bar = 48 nm. (C) Flowchart of image processing and reconstruction.

**Figure 1-supplement 2.**
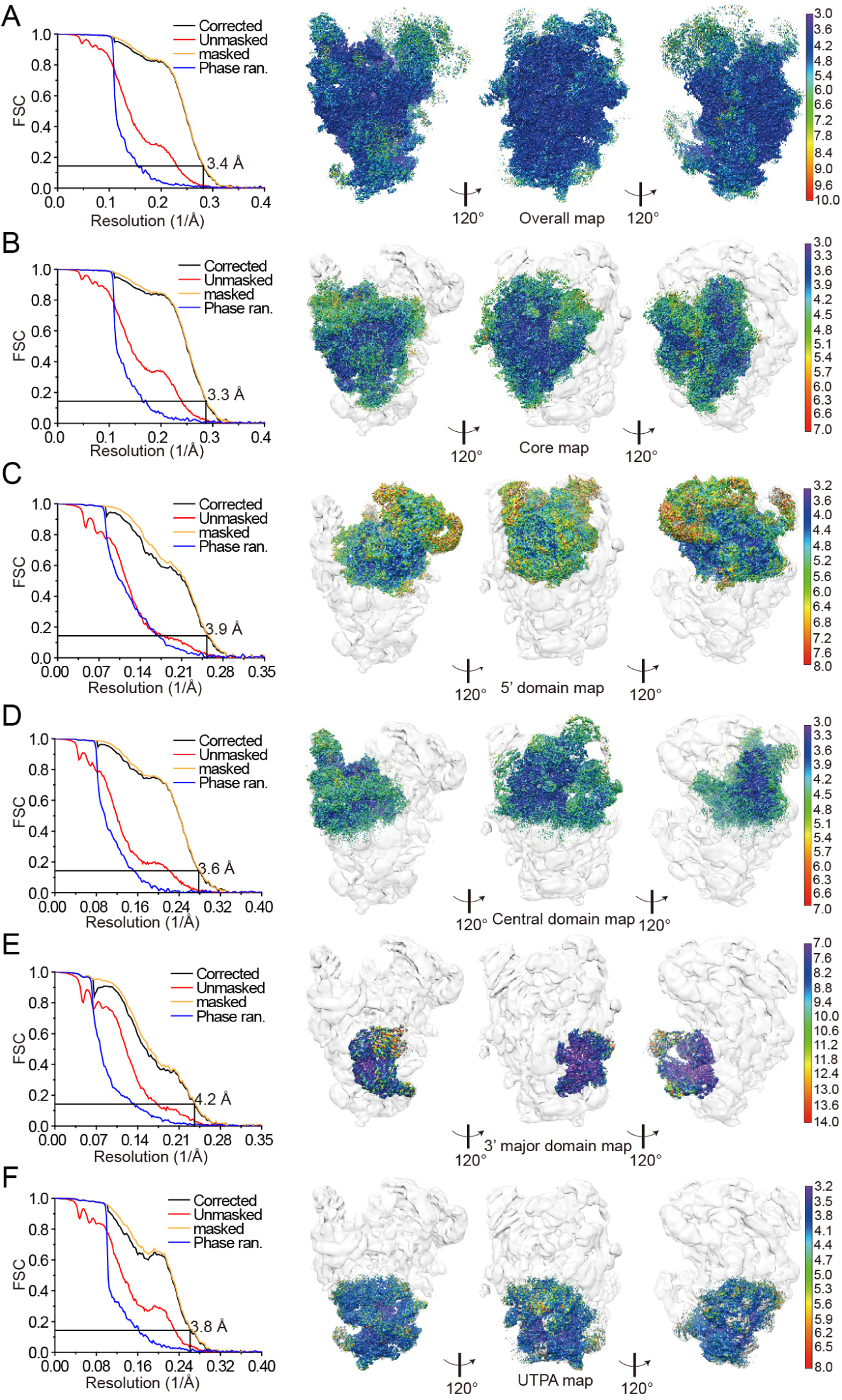
Resolution estimation of the overall and focused cryo-EM maps of Mtr4-depleted 90S. (A) The overall map. (B) The core map. (C) The 5’ domain map. (D) The central domain map. (E) The 3’ major domain map. (F) The UTPA map. Fourier shell correlation (FSC) curves of the unmasked, masked, phase-randomized and corrected map are represented by red, green, blue and black lines, respectively. Each map is shown in three views related by 120 degree rotation and colored by local resolution values. The focused maps in B-E are displayed on a transparent overall map low-pass filtered to 20 Å.

**Figure 1-supplement 3.**
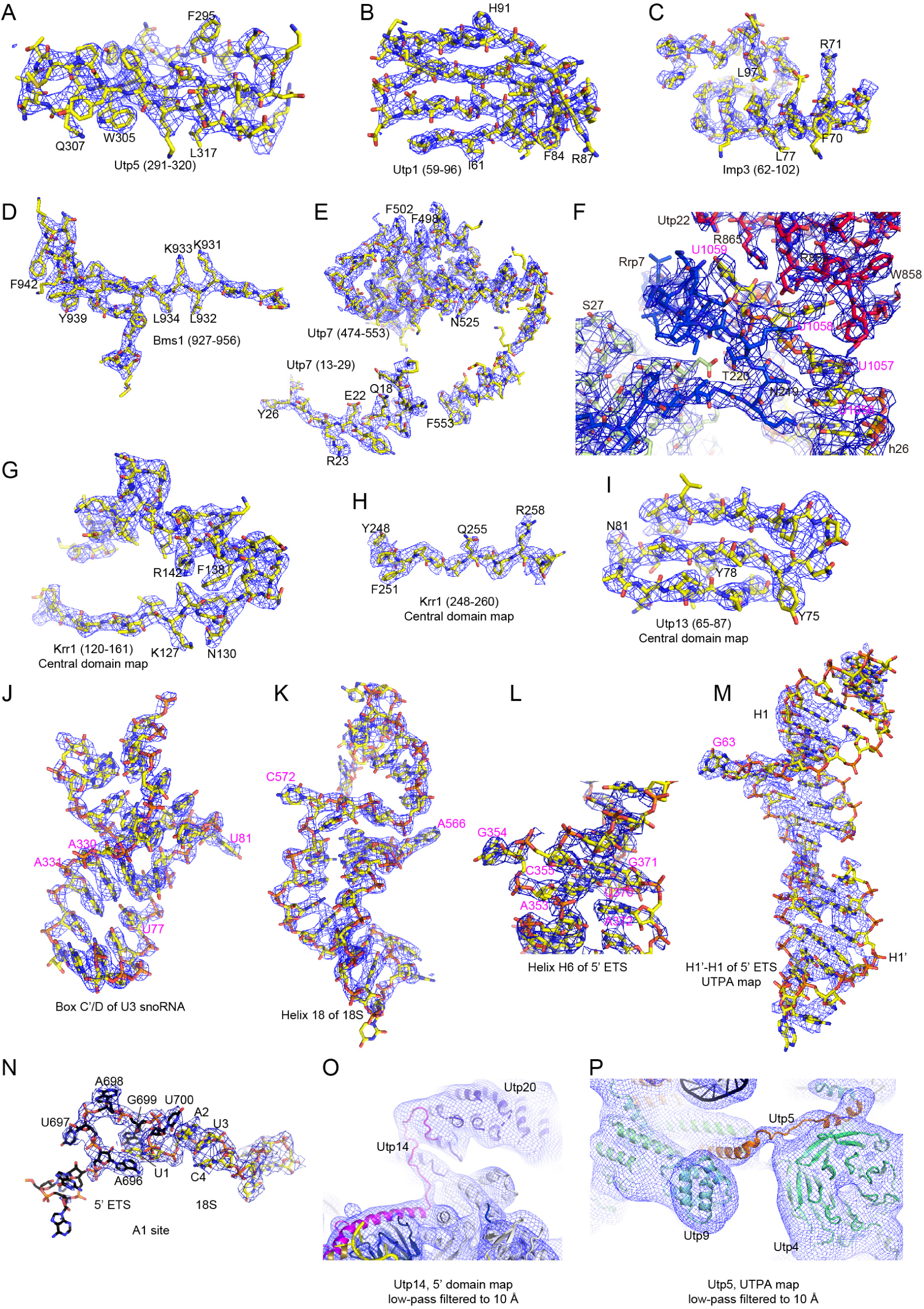
Representative cryo-EM density maps with the fitted structure model. (A) Utp5. Selected residues with clear side chain density are labeled. The overall map is displayed unless stated otherwise. (B) Utp1. (C) Imp3. (D) Bms1. (E) Utp7. (F) The tip of helix 26 of 18S rRNA and its binding proteins. This view is the same as Figure 3C. (G) The KH domain of Krr1 on the central domain map. (H) The C-terminal tail of Krr1 on the central domain map. (I) Utp13 on the central domain map. (J) Box C’/D of U3 snoRNA. (K) Helix 18 of 18S rRNA. (L) H6 of 5’ ETS. This view is the same as Figure 6F. (M) Helices H1’ and H1 of 5’ ETS on the UTPA map. (N) The A1 site separating 5’ ETS and 18S rRNA. (O) The C-terminal tail of Utp14 on the 5’ domain map low-pass filtered to 10 Å. (P) The C-terminal tail of Utp5 on the UTPA map low-pass filtered to 10 Å.

**Figure 1-supplement 4.**
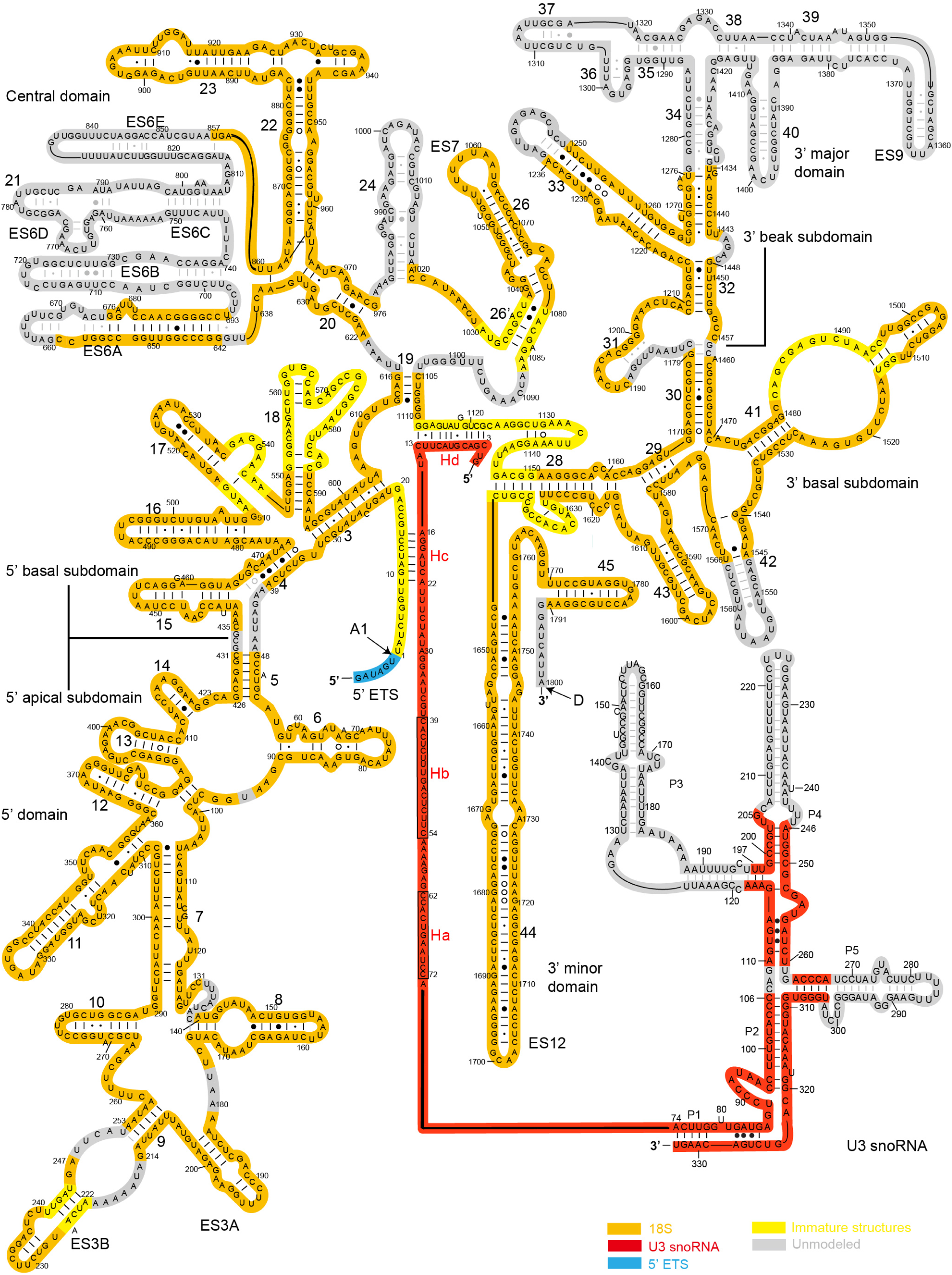
Secondary structure model of 18S rRNA and U3 snoRNA in Mtr4-depleted 90S preribosome. The sequences of 18S rRNA that adopt a similar or different secondary structure as in the mature SSU are shaded in orange or yellow, respectively. U3 snoRNA and a short segment of 5’ ETS are shaded in red and blue. The unmodeled regions are shaded in grey. The A1 and D processing sites, name of RNA helices are indicated.

**Figure 1-supplement 5.**
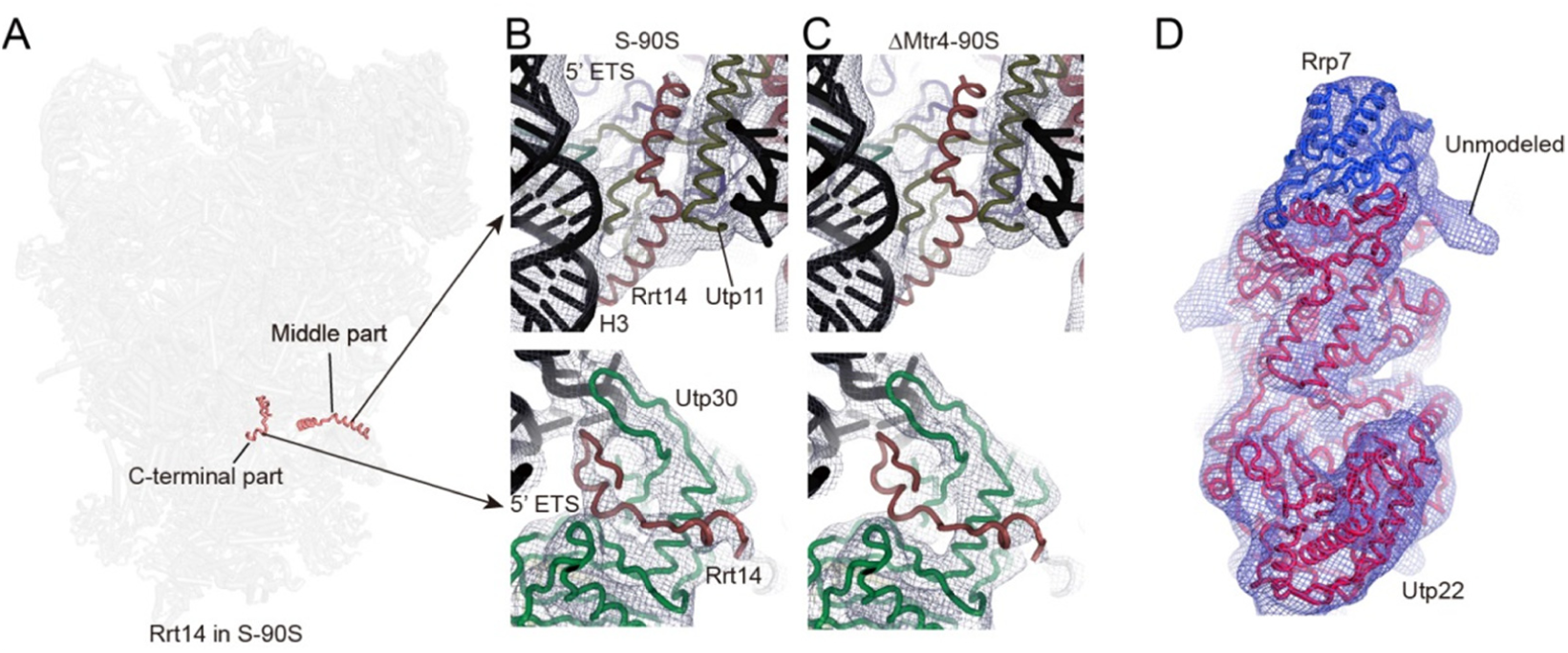
Rrt14 and Rrp5 display extremely weak densities in Mtr4-depleted 90S. (A) Location of Rrt14 in the S-90S structure. (B-C) Cryo-EM density of Rrt14 in S-90S (B) and Mtr4-depleted 90S (C) low-pass filtered to 8 Å (D) Cryo-EM density of Mtr4-depleted 90S low-pass filtered to 10 Å showing an unmodeled weak density attached to Utp22 at the location of the TRP domain of Rrp5.

**Figure 5-figure supplement 1.**
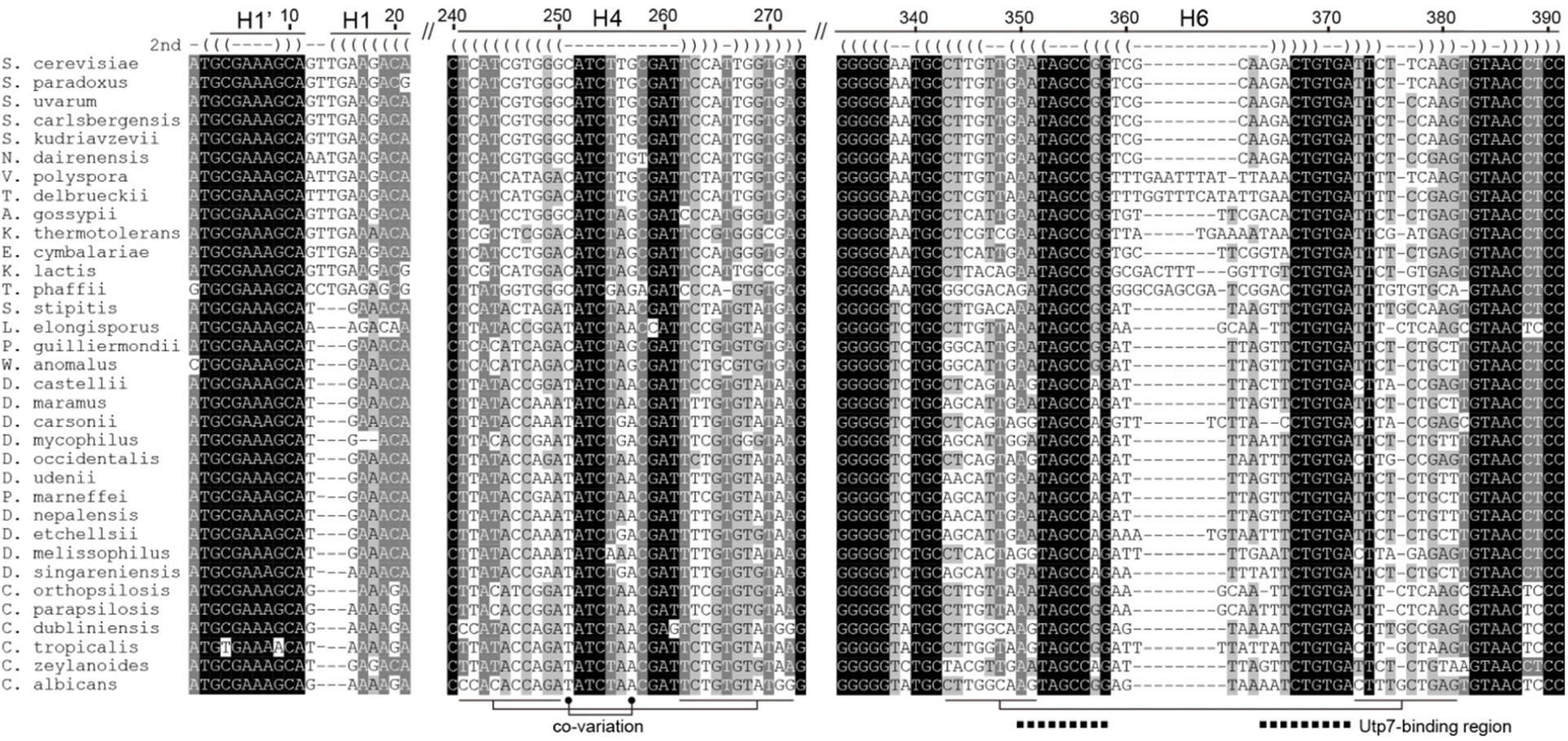
Multiple sequence alignment of the 5’ ETS from Saccharomycotina. 34 sequences of 5’ ETS were aligned. Only the H1’, H4 and H6 region are displayed. Residues that have >95%, 95-80% and 80-60% identity are shaded with black, grey and light grey, respectively. Base-paired residues are annotated with parentheses on the top. Positions showing strong co-variation are connected at the bottom. The nucleotides of H6 that bind Utp7 are marked with squares.

**Supplementary Table 1.** Statistics of data collection, structural refinement and model validation

|  |  |  |  |  |  |  |
| --- | --- | --- | --- | --- | --- | --- |
| **Data collection** |  |  |  |  |  |  |
| EM equipment | FEI Titan Krios |  |  |  |  |  |
| Voltage (kV) | 300 |  |  |  |  |  |
| Detector | K2 Summit |  |  |  |  |  |
| Grid | Quantifoil 1.2/1.3 |  |  |  |  |  |
| Micrographs | 4,723 |  |  |  |  |  |
| Particles for 3D classification | 362,333 |  |  |  |  |  |
| Pixel size (Å) | 1.06 |  |  |  |  |  |
| Defocus range (µm) | 1.5-2.5 |  |  |  |  |  |
| Electron dose (e<sup>-</sup>/Å<sup>2</sup>) | 50 |  |  |  |  |  |
| **Map refinement** | Overall map | Core map | 5' domain map | Central domain map | 3' major domain map | UTPA map |
| Particles for refinement | 128,637 | 121,139 | 75,499 | 95,349 | 83,840 | 120,456 |
| Overall resolution of map (Å) | 3.4 | 3.3 | 3.9 | 3.6 | 4.2 | 3.8 |
| Map sharpening B-factor (Å<sup>2</sup>) | -100 | -99 | -125 | -98 | -158 | -136 |
| **Model composition** |  |  |  |  |  |  |
| Protein chains | 68 |  |  |  |  |  |
| Protein residues | 24,567 |  |  |  |  |  |
| RNA chains | 3 |  |  |  |  |  |
| RNA bases | 2008 |  |  |  |  |  |
| **Structural refinement** |  |  |  |  |  |  |
| CC_mask | 0.767 |  |  |  |  |  |
| CC_volume | 0.744 |  |  |  |  |  |
| RMSD bonds (Å) | 0.011 |  |  |  |  |  |
| RMSD angles (°) | 1.132 |  |  |  |  |  |
| **Validation (protein)** |  |  |  |  |  |  |
| MolProbity score | 2.02 |  |  |  |  |  |
| Clashscore | 7.09 |  |  |  |  |  |
| Good rotamers (%) | 98.74 |  |  |  |  |  |
| Ramachandran plot favored (%) | 89.91 |  |  |  |  |  |
| Ramachandran plot allowed (%) | 9.99 |  |  |  |  |  |
| Ramachandran plot outliers (%) | 0.10 |  |  |  |  |  |

**Supplementary Table 2.**
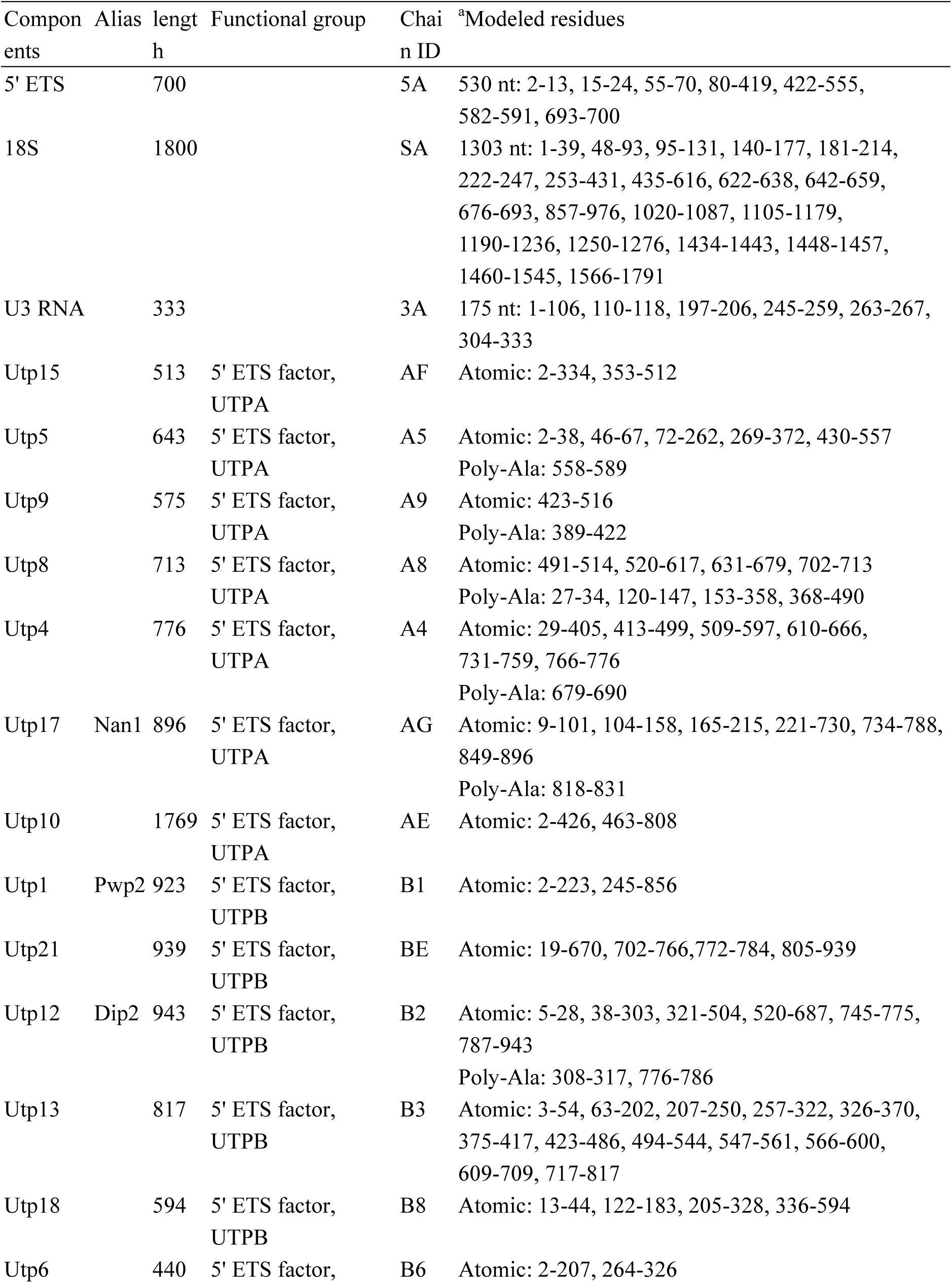
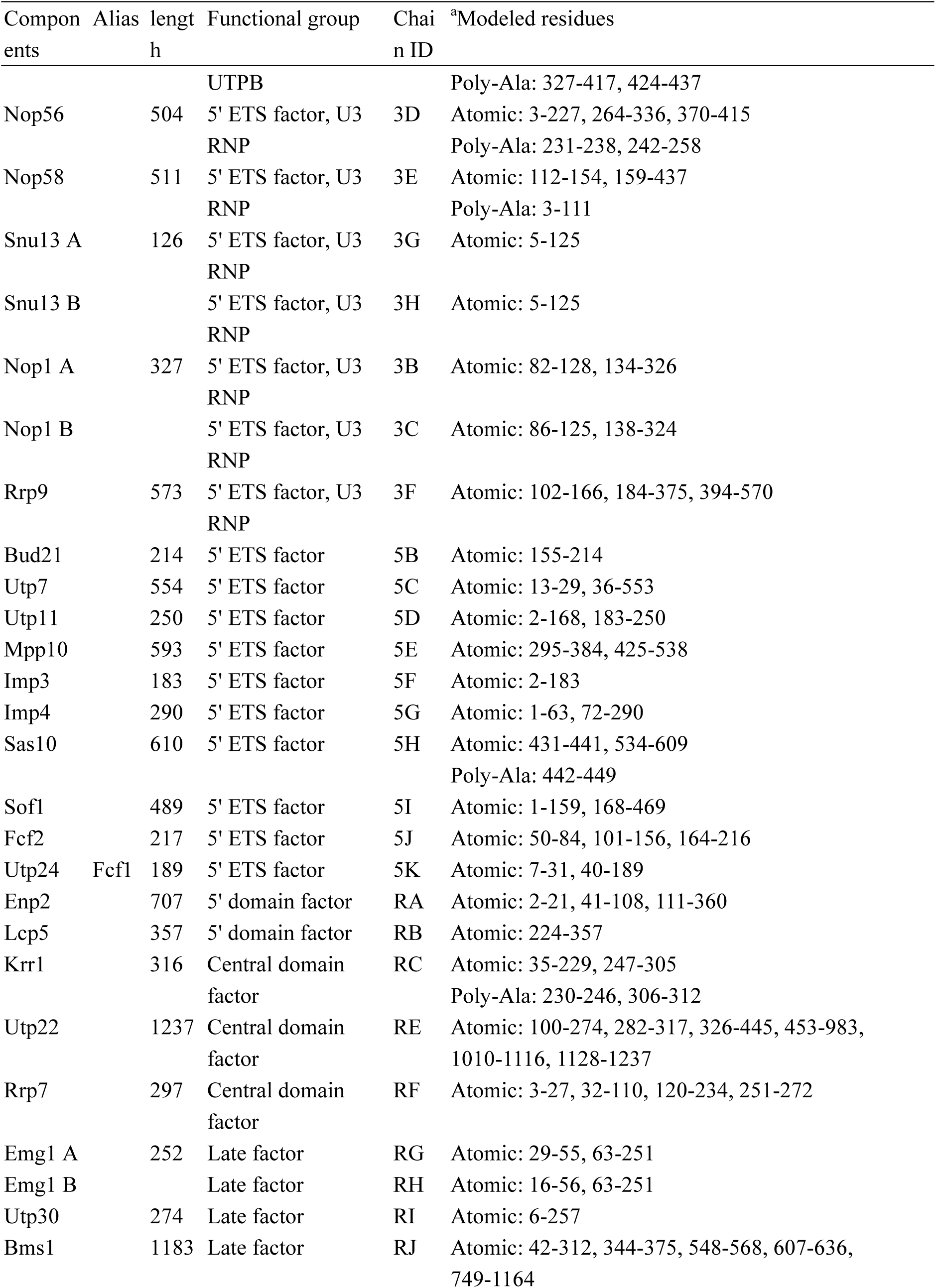
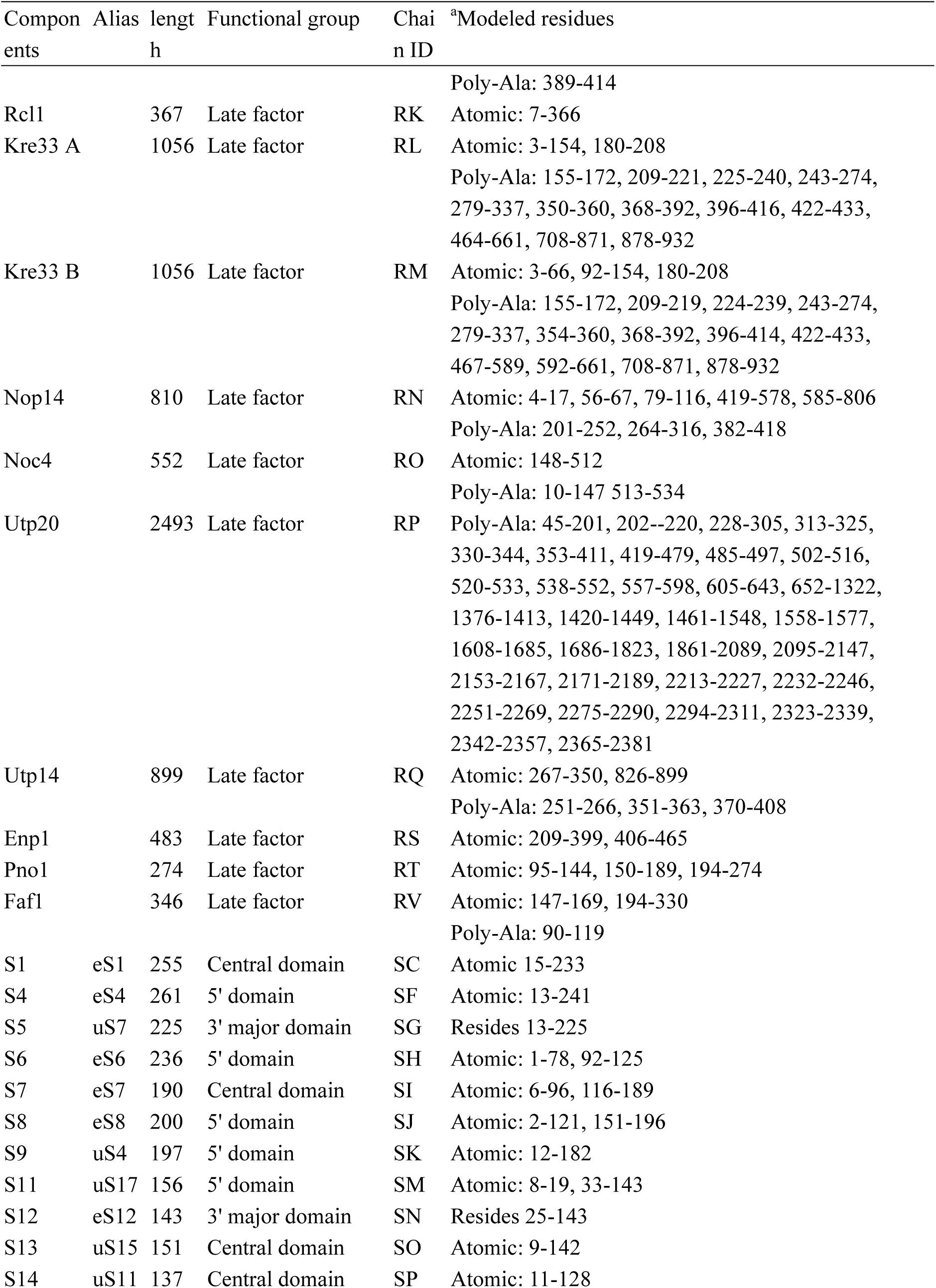
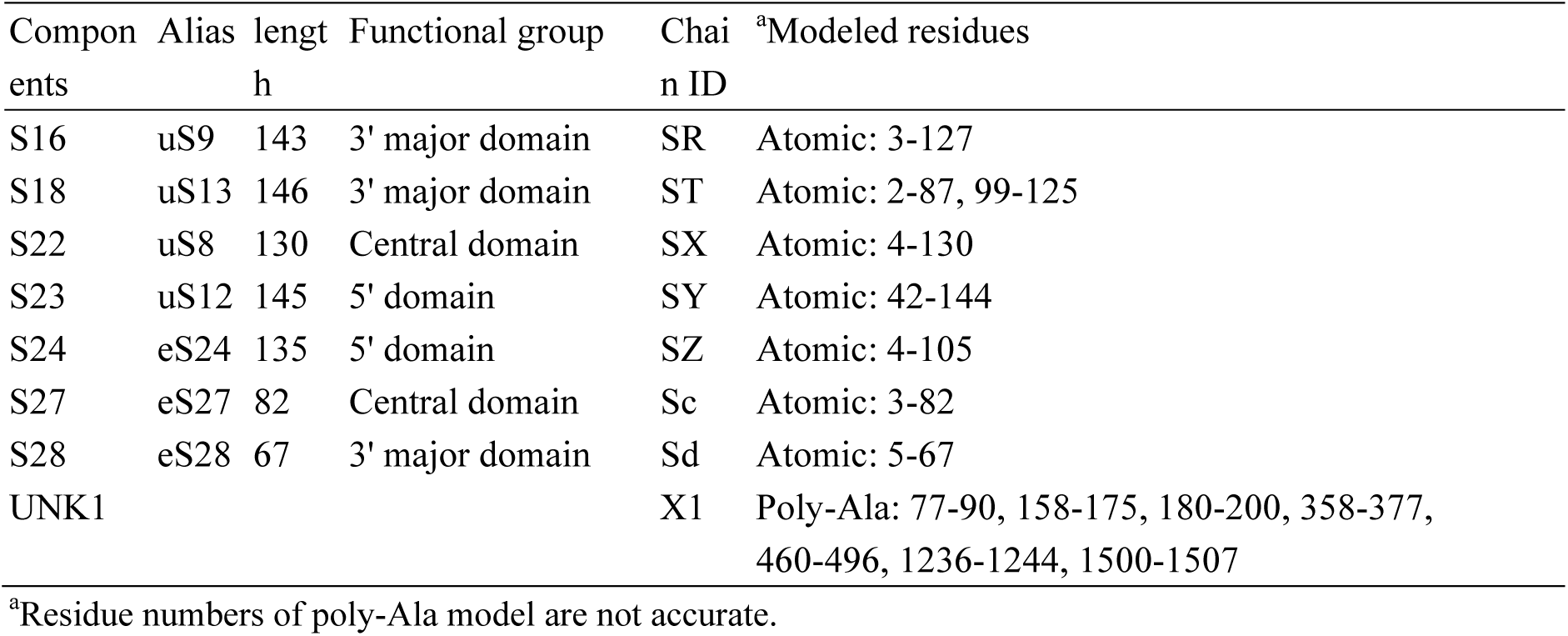
Composition of Mtr4-depleted 90S structure.

